# Homeostatic Plasticity Requires Remodeling of the Homer-Shank Interactome

**DOI:** 10.1101/2020.03.26.010314

**Authors:** Whitney E. Heavner, Haley Speed, Jonathan D. Lautz, Edward P. Gniffke, Karen B. Immendorf, John P. Welsh, Stephen E.P. Smith

## Abstract

Neurons maintain constant levels of excitability using homeostatic scaling, which adjusts relative synaptic strength in response to large changes in overall activity. It is still unknown how homeostatic scaling affects network-level protein interactions in the synapse despite extensive reporting of individual scaling-associated transcriptomic and proteomic changes. Here, we assessed a glutamatergic synapse protein interaction network (PIN) composed of 380 binary interactions among 21 protein members to identify protein complexes altered by synaptic scaling in vitro and in vivo. In cultured cortical neurons, we observed widespread bidirectional PIN alterations during up- and downscaling that reflected rapid glutamate receptor shuttling via synaptic scaffold remodeling. Sensory deprivation of the barrel cortex caused a PIN response that reflected changes in mGluR tone and NMDAR-dependent metaplasticity, consistent with emerging models of homeostatic plasticity in the barrel cortex that restore excitatory/inhibitory balance. Mice lacking *Homer1* or *Shank3B* did not undergo normal PIN rearrangements, suggesting that these Autism Spectrum Disorder (ASD)-linked proteins serve as structural hubs for synaptic homeostasis. Our approach demonstrates how changes in the protein content of synapses during homeostatic plasticity translate into functional PIN alterations that mediate changes in neuron excitability.

## Introduction

Neurons maintain constant relative cell-wide synaptic strengths, despite varying network activity, through a non-Hebbian form of neural plasticity called homeostatic synaptic scaling (*1*). At the molecular level, homeostatic scaling adjusts the number of postsynaptic glutamate receptors up or down to compensate for prolonged decreased or increased synaptic activity, without compromising the distributed information content of each synapse (*2*–*7*). In addition to, or in cooperation with, other homeostatic mechanisms, such as intrinsic homeostatic plasticity (*8*) and sliding-scale plasticity (also called “metaplasticity”) (*9*), synaptic scaling in vivo restores the activity of a neuron to an initial set point after prolonged sensory deprivation (*10*, *11*) or prolonged activation (*12*) (reviewed in (*13*)). Homeostatic plasticity is thought to be crucial for preventing synaptic saturation, thus allowing for new memory formation, and may be disrupted in several neurological disorders, including Alzheimer’s disease (*14*) and ASD (*15*–*18*).

Prior studies have catalogued extensive molecular alterations associated with homeostatic scaling. At the most basic level, upscaling is mediated by insertion of GluR2-containing AMPA receptors and alterations in AMPA receptor co-association with synaptic scaffolds (*19*). Additional critical molecular pathways have been evaluated using single candidate-based approaches. For example, fluctuations in intracellular Ca^2+^ levels lead to CaMKIV activation, which drives changes in gene expression (*20*) and downstream synaptic accumulation of trafficking proteins, such as GRIP1 (*21*). Activity blockade also de-represses retinoic acid synthesis, leading to increased local translation and synaptic insertion of AMPA receptors (*22*). In addition, widespread changes in protein ubiquitination have been documented during scaling (*23*), and omics-scale studies have catalogued extensive alterations to the transcriptome (*20*, *24*–*26*) and proteome (*27*, *28*). These changes in transcript and protein abundance are often bidirectional, or opposite, for upscaling versus downscaling. At the network level, binding of the immediate early gene *Homer1a* to the metabotropic glutamate receptor mGluR5 (*4*, *29*) and regulation of the expression of the scaffolding protein/transcription factor *Ctnnb1* by the chromatin reader L3MBTL1 (*24*) play important roles. Taken together, these studies highlight a complex molecular process through which changes in synaptic activity lead to compensatory changes in synaptic excitability.

Just as mRNA levels do not predict protein expression in complex systems (*30*, *31*), protein levels *alone* do not determine physiological outcomes. The interactions between newly synthesized or degraded molecules and the existing protein complexes in the cell determine the physiological effects of altered protein turnover. In addition to absolute protein abundance, post-translational modifications and the presence or absence of functional binding partners influence protein complex composition. In fact, the combinatorial complexity of the protein interactome, as well as its incomplete characterization (*32*), make predicting the physiological effect of altered protein abundance exceedingly difficult. Here, by quantitatively measuring changes to the macromolecular protein complexes that control the localization of glutamate receptors at the postsynapse, we bridge the gap between well-characterized protein abundance changes and the electrophysiological effects that occur during homeostatic scaling. Using an emerging proteomic technique, <u>Q</u>uantitative <u>M</u>ultiplex co-<u>I</u>mmunoprecipitation (QMI), we characterize changes to a synaptic PIN consisting of 21 unique synaptic proteins (380 binary interactions) following upscaling and downscaling and demonstrate how altered protein production and degradation result in widespread, coordinated changes to the functional PINs that mediate synaptic excitability. Additionally, using sensory deprivation in two different mouse models of ASD, we demonstrate that the synaptic scaffolding proteins Homer1 and Shank3 are required for PIN rearrangements characteristic of homeostatic plasticity in vivo.

## Results

### Upscaling and downscaling induce distinct changes in protein interaction networks

We cultured dissociated cortical neurons from postnatal day (P) 0 wild-type mice for 15-19 days, then treated each culture for 48 hours with tetrodotoxin (TTX, 2μM) to induce upscaling, bicuculine (BIC, 40μM) to induce downscaling, or dimethylsulfoxide (DMSO, 0.4%), a vehicle control. We confirmed upscaling and downscaling using surface expression of GluR1 (*33*), which was significantly elevated after TTX treatment and reduced after BIC treatment (fold-change=2.32 and 0.256, *P*-value=0.04 and 0.01, respectively) (**fig. 1A**). Previous studies have reported widespread changes in the levels of synaptic proteins after 2 to 72 hours of upscaling or downscaling in vitro (*23*, *27*, *28*), so we quantified total levels of several synaptic proteins by western blot after 24 and 48 hours of scaling (**fig. 1B, C**). After 24 hours of downscaling, we observed a significant decrease in PSD95 (*P*=0.03), Fyn (*P*=0.005), and the glutamate receptors GluR1 (*P*=0.003) and mGluR5 (*P*=0.01), consistent with previous reports (*23*, *34*) (*27*). By contrast, after 24 hours of upscaling, only mGluR5 changed significantly (*P*=0.005), and was increased, opposite downscaling, Similarly, GluR1 tended to increase but was not statistically significant (*P*=0.1), suggesting that protein abundance changes are not completely bidirectional. By 48 hours of upscaling, GluR1 had significantly increased (*P*=0.03), while mGluR5 levels appeared to have normalized, and Homer1 was significantly reduced (*P*=0.03). By contrast, mGluR5, PSD95, and Fyn remained reduced after 48 hours of downscaling (*P*=0.009, 0.007, 0.04, respectively), while GluR1 levels appeared to have normalized, and Homer1 was slightly but significantly reduced (*P*=0.05), again demonstrating that protein abundance changes are not exactly opposite for upscaling and downscaling.

**Fig. 1.**
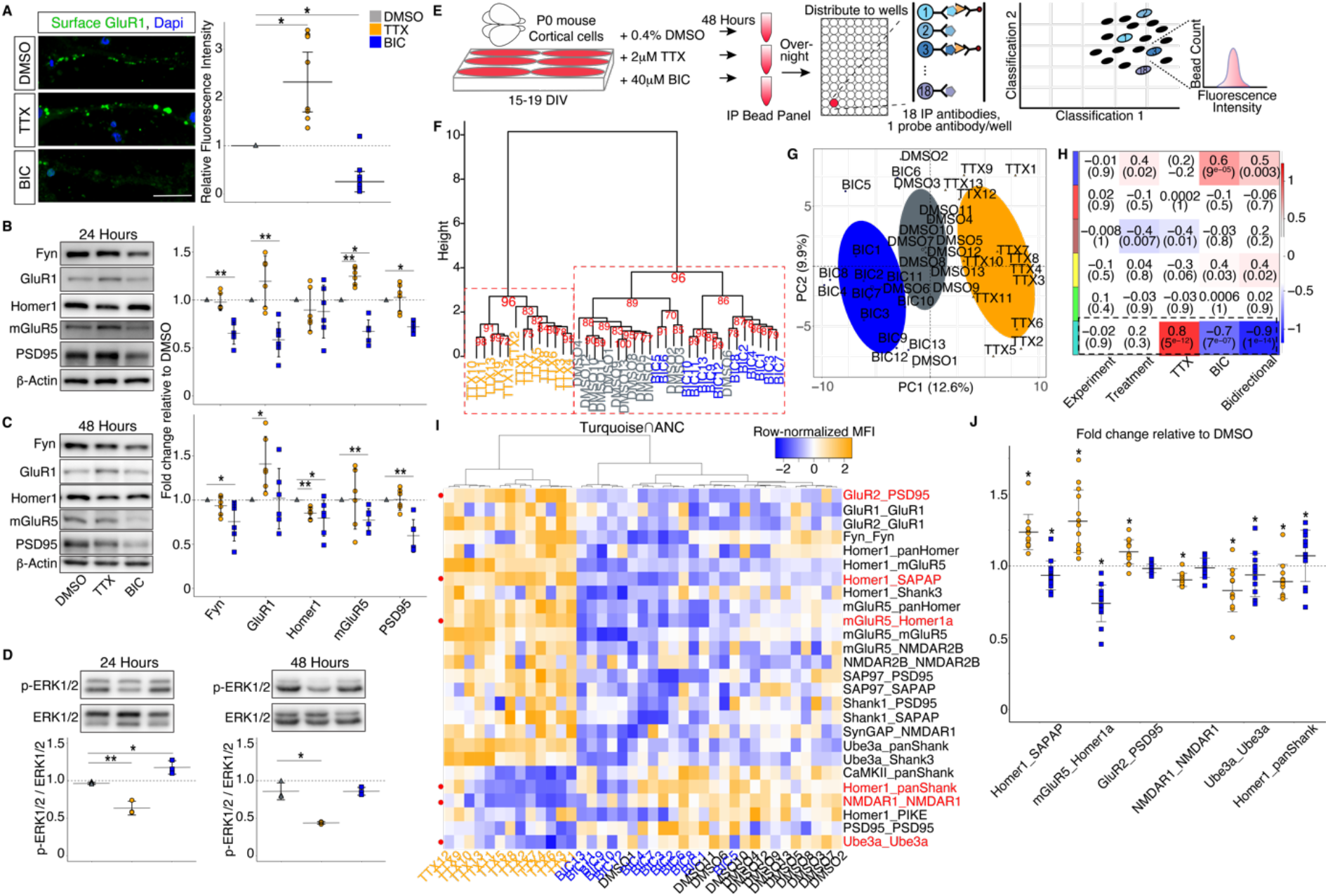
Prolonged increased or decreased activity of cultured cortical neurons causes network-level changes to multiprotein complexes. **(A)** Surface GluR1 (green) and Dapi (nuclei, blue) in cortical neurons treated with DMSO (control), TTX, or BIC. Mean fluorescence intensity of GluR1 relative to control is shown in the scatter plot to the right (N=9 cells from 3 biological replicates; DMSO = grey throughout, TTX = orange throughout, BIC = blue throughout). **(B,C)** Representative western blots of total levels of select synaptic proteins after 24 (B) or 48 (C) hours of DMSO, TTX, or BIC treatment. Mean protein levels relative to control after treatment are shown in the scatter plot to the right (N=6 biological replicates). **(D)** Representative western blots of total levels of ERK1/2 and p-ERK1/2 after 24 or 48 hours of treatment. Mean ratios of p-ERK to total ERK are shown in the scatter plots below (N=3 biological replicates). **(E)** Workflow for QMI analysis of in vitro homeostatic scaling. **(F)** Hierarchical clustering (AU *P*-values shown in red) and **(G)** principal component analysis of 39 samples representing 3 different treatment conditions (13 biological replicates per condition, 48 hours of treatment) using MFI values. **(H)** Module-trait relationship heatmap showing the correlation (top number) and *P*-value for each module-trait pair (N=13 biological replicates). **(I)** Heatmap of row-normalized MFIs of all ANC-significant PiSCES in the turquoise module. Samples on the X-axis are arranged by unsupervised hierarchical clustering. PiSCES on they Y-axis are arranged into those that go up with TTX/down with BIC and vice versa. **(J)** Scatter plots of the mean log_2_(fold change) of select PiSCES after 48 hours of treatment for each set. PiSCES in red in I are represented by scatter plot in J. Scale bar in A=20μm. Error bars represent SD. For A-D: (*) *P*-value < 0.05; (**) *P*-value < 0.01. For J: (*) ANC-significant.

Prolonged changes in neural activity have been shown to affect the MEK-ERK pathway, leading to bidirectional changes in ERK1/2 phosphorylation upstream of changes to activity-dependent mRNA and protein production (*35*). We measured the relative levels of phosphorylated (p-) ERK1/2 after 24 and 48 hours of scaling **(fig. 1D)** and found that upscaling was associated with a significant decrease in the ratio of p-ERK1/2 to total ERK1/2 after 24 hours (*P*=0.004), which remained significantly decreased after 48 hours (*P*=0.02). Conversely, downscaling was associated with an increased ratio of p-ERK1/2 to total ERK1/2 after 24 hours (*P*=0.02), which normalized after 48 hours, consistent a previous report showing negative feedback onto p-ERK1/2 through changes to surface glutamate receptors (*35*, *36*). Together with the above data, these results demonstrate that prolonged activity perturbation alters surface GluR1, synaptic protein abundance, and ERK1/2 activation consistent with prevailing views of homeostatic scaling.

To determine how homeostatic scaling affects PINs, we next evaluated changes in synaptic protein complexes after 48 hours of upscaling or downscaling using quantitative multiplex immunoprecipitation (QMI). QMI takes advantage of fluorophore-embedded microspheres to immunoprecipitate multiple protein targets from a single sample simultaneously (*37*, *38*). Biotinylated probe antibodies detect selected co-associated proteins, and, following streptavidin-PE secondary labeling, the median fluorescence intensity (MFI) of each IP-probe combination is measured on a flow cytometer and averaged across technical replicates (see methods) **(fig. 1E)**. We refer to multiprotein complexes detected by QMI as “<u>P</u>roteins in <u>S</u>hared <u>C</u>omplexes detected by surface <u>E</u>pitope<u>s</u>” (PiSCES), to highlight the fact that the measured interactions are not necessarily direct. The QMI panel described in this report contains 21 validated synapse-associated proteins (*37*, *39*) and measures 380 binary PiSCES for each experiment **(table 1).**

**Table 1.**
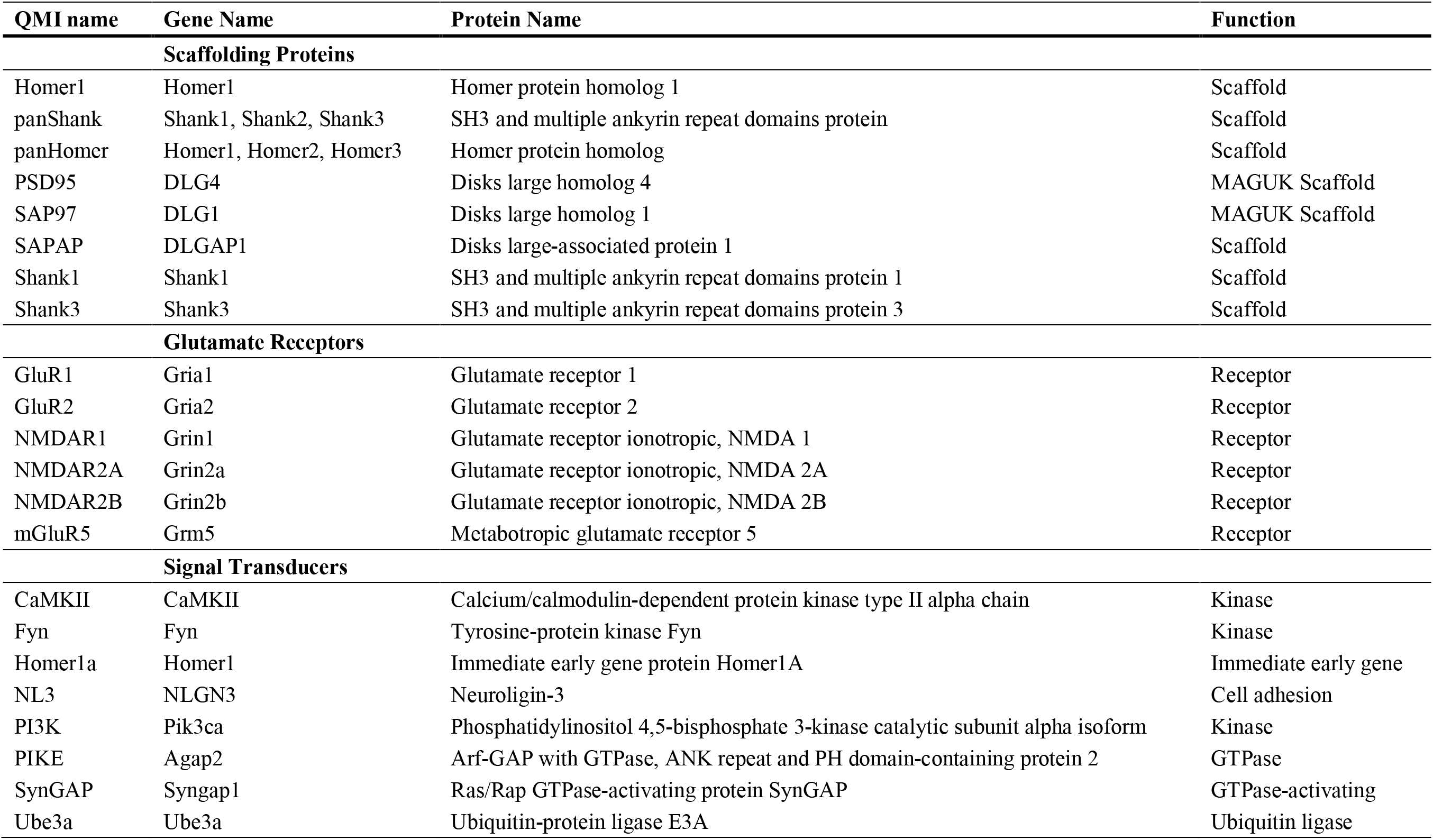
Gene names, proteins names, and functions of protein targets that comprise the QMI assay.

Network-level differences in PiSCES between TTX, BIC, and DMSO-treated cells (N=13 biological replicates) were evaluated using hierarchical clustering and principal component analysis (PCA). Approximately unbiased (AU) *P*-values (*40*), determined by multiscale bootstrap resampling, identified two major clusters correlated with treatment: 1) all TTX samples formed one cluster, while 2) BIC samples clustered with DMSO; however, 11 of the 13 BIC samples formed their own subcluster (**fig. 1F**). PCA showed separation of the three conditions along principal component (PC) 1 (**fig. 1G**). These results demonstrate that there is more similarity within a treatment group than between groups and imply that prolonged increased or decreased activity causes distinct network-level changes in synaptic PINs, in addition to changes in protein levels.

### Upscaling and downscaling induce bidirectional scaffold changes in vitro

As previously described (*38*), QMI relies on a combination of two statistical tests: 1) <u>a</u>daptive <u>n</u>onparametric analysis with an empirical alpha <u>c</u>utoff (ANC) and 2) weighted <u>c</u>orrelation <u>n</u>etwork <u>a</u>nalysis (CNA) (*41*). CNA identified one primary module (“turquoise”) that was positively correlated with TTX (*P*=5×10^−12^) and negatively correlated with BIC (*P*=7×10^−7^) (**fig. 1H**). This module was merged with ANC-significant PiSCES (see methods) to identify 26 high-confidence interactions that met the criteria for both independent statistical approaches. When these ANC∩CNA PiSCES were plotted as a heatmap and samples were ordered using unsupervised clustering, samples clustered by treatment type, and a clear pattern emerged in which PiSCES changed bidirectionally between TTX and BIC treatment, while DMSO served as an intermediate (**fig. 1I**). The mean log_2_ fold changes of selected PiSCES are shown in (**fig. 1J**).

PiSCES changes in the turquoise module broadly reflected shuttling of AMPA and mGluR receptors throughout the membrane: interactions between GluR1 and GluR2 (expressed as GluR1_GluR2) and GluR2_PSD95 increased with TTX and decreased with BIC, while mGluR5_Homer1 and mGluR5_Homer1a followed a similar pattern (**fig. 1I, J**). These data are consistent with upregulation of mGluR and AMPAR signaling during upscaling and downregulation during downscaling (*4*, *19*, *22*) and highlight an underappreciated role for mGluRs in homeostatic scaling. In fact, Homer1_mGluR5 was the single largest log_2_ fold change we detected, and mGluR interaction with Homer1a, which is critical for mediating homeostatic plasticity during sleep (*29*), has been shown to affect its axonal and dendritic targeting and interaction with the Homer/Shank scaffold (*42*, *43*).

In fact, bidirectional scaffold protein rearrangements were prevalent in the turquoise module: PSD95_SAP97, SAP97_ SAPAP, Homer1_SAPAP, Shank1_ SAPAP, and Shank1_PSD95 were increased with TTX/decreased with BIC, reflecting rearrangement of the scaffold with DLG family proteins. Conversely, Homer1_panShank, and PiSCES involving Shank1, panShank, and CaMKII, which regulates the turnover of AMPARs during scaling via phosphorylation of SAPAP (*44*), decreased with TTX and increased with BIC (**fig. 1I, J**). Compared with protein levels changes, changes in protein associations between scaffold proteins and glutamate receptors appeared to be similarly, if not more broadly, bidirectional. A few PiSCES reflecting self-associations, however, such as NMDAR1_NMDAR1 and Ube3a_Ube3a, decreased with both TTX and BIC (**fig. 1J**). Collectively, these data suggest that homeostatic synaptic scaling tunes synaptic strength through bidirectional changes in associations between synaptic scaffold proteins and glutamate receptors.

Three modules were only marginally correlated with treatment (“brown,” “blue,” and “yellow”) (**fig. 1H**). Each contained additional ANC-significant PiSCES, but did not segregate by treatment when clustered using only PiSCES in the given module **(fig. S2)**. Therefore, while we hypothesize that these PiSCES represent biologically-relevant modules of interacting proteins that co-vary in cultured cells, they seemed to be unrelated to TTX/BIC treatment and were excluded from further analysis. Taken together, these results identify a high-confidence set of multiprotein complexes that rearrange bidirectionally during 48 hours of homeostatic scaling.

### Protein interaction changes broadly stabilize by 12 hours of scaling

Different sets of proteins are newly synthesized after prolonged activity perturbation for a time period lasting at least 2-24 hours (*27*, *28*). We therefore asked whether different PiSCES changed at different timepoints over a 48-hour period of upscaling or downscaling. We performed QMI after 12 hours (N=4) and 24 hours (N=4) of TTX and BIC treatment and compared PiSCES changes between all three time points (using 48 hours set 2 from above). Unsupervised hierarchical clustering of all interactions (MFI>100) separated TTX, BIC, and DMSO conditions into discrete groups (**fig. 2A**). AU *P*-values identified three major clusters correlated with treatment: 1) all TTX samples formed one cluster, 2) BIC 48-hours formed an independent cluster, and 3) BIC 12-hours and BIC 24-hours clustered with DMSO. Similarly, PCA showed that all three TTX groups separated from DMSO control groups along PC1 (**fig. 2B**). By contrast, the BIC 12-hour group overlapped with the DMSO groups, but the BIC 24- and 48-hour groups separated from DMSO along PC1 in the opposite direction of TTX.

**Fig. 2.**
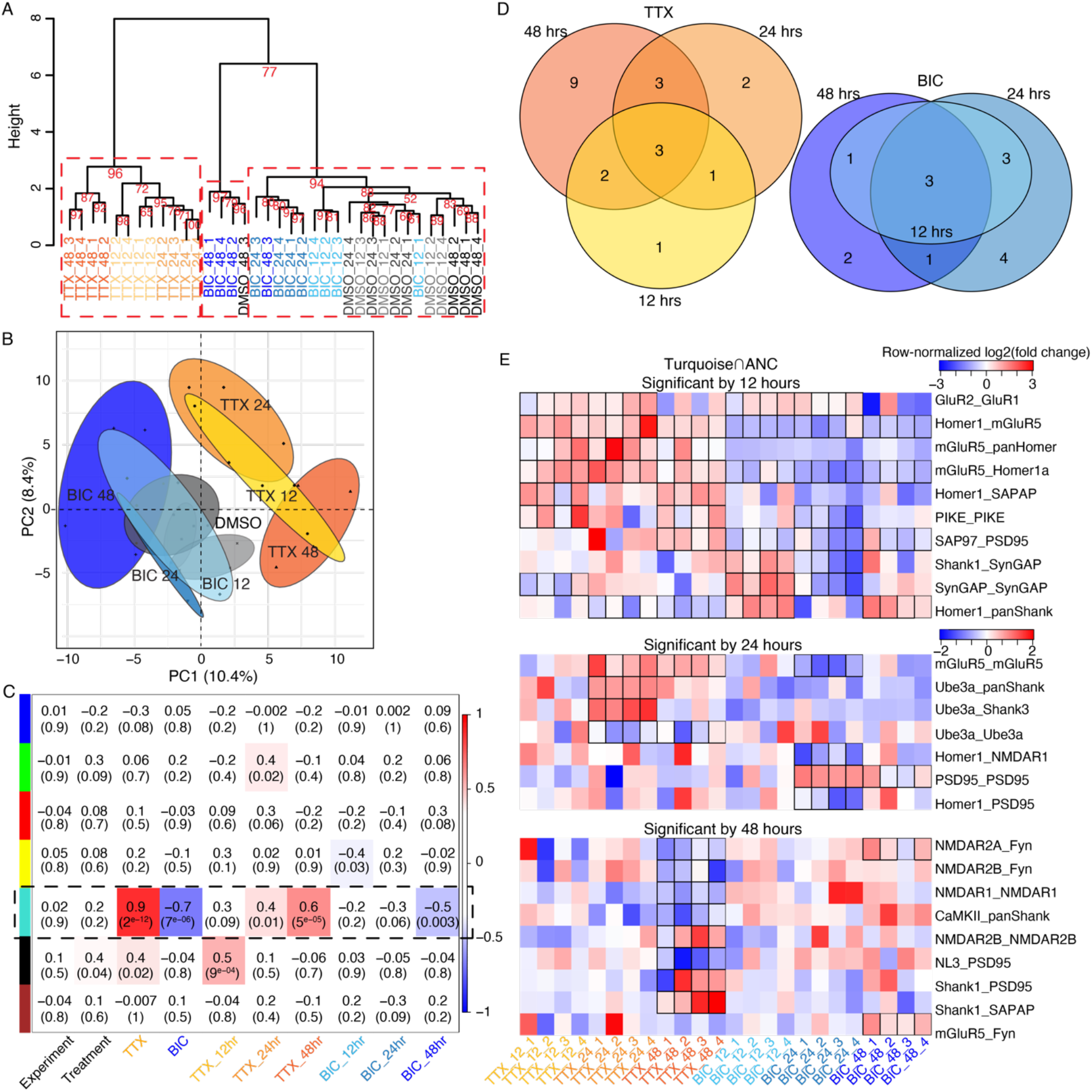
Many PiSCES changes stabilize within 12 hours of homeostatic scaling. **(A)** Hierarchical clustering of three sets of experiments (N=4 biological replicates per set per condition) covering three time points (12 hours, 24 hours, and 48 hours) shows separation of all 12 TTX samples into one significant cluster, regardless of time, while BIC samples form distinct sub-clusters. AU *P*-values are shown in red. **(B)** PCA of all PiSCES measurements (MFI>100) shows separation of TTX and BIC conditions along PC1. **(C)** Module-trait relationship heatmap showing the correlation (top number) and *P*-value for each module-trait pair. **(D)** Venn diagrams showing the overlap of ANC-significant PiSCES changes at each timepoint. **(E)** Row-normalized heatmaps of the mean log_2_(fold change) after treatment for every ANC-significant PiSCES in the turquoise module. DMSO is set to 0 (not shown), so that blues indicate decreased interaction, and reds indicate increased interaction. Outlines indicate sets for which the PiSCES measurement was significantly different from DMSO. PiSCES are arranged into groups according to the earliest time point at which they became significant.

We next used CNA as described above to identify high-confidence PiSCES that changed together in response to treatment. We identified one major bidirectional module similar to the turquoise module described above (**fig. 2C**). When we asked if any timepoint alone was better correlated with this module compared to treatment as a single trait independent of time; we found that the 12-hour timepoint showed no correlation, the 24-hour timepoint showed some correlation but was only significant for TTX, and the 48-hour timepoint showed strong correlation for both TTX and BIC, although neither was as strongly correlated with the bidirectional module as TTX or BIC independent of time. Collectively, these data demonstrate that TTX and BIC induce consistent, bidirectional PiSCES changes that increase in magnitude over 48 hours.

To identify which PiSCES contributed to the bidirectional module, we merged ANC hits at each timepoint with the module members (**fig. 2D,E**). At 12 hours, 7 PiSCES were identified for both TTX and BIC, all of which were significant in at least one other timepoint. For TTX, with the exception of SynGAP_SynGAP, all of these PiSCES were elevated, including GluR2_GluR1 and Homer1_mGluR5, which remained elevated throughout the time course (**fig. 2E**), reflecting rapid and persistent AMPAR insertion and remodeling of the Homer scaffold, which adjusts mGluR tone, during the initial stages of upscaling. Conversely, the Homer1-containing PiSCES that were significantly increased during upscaling were decreased during downscaling throughout the time course, which, in addition to the sustained reduction in SAP97_PSD95, reflected early and persistent synapse weakening by 12 hours of elevated activity (*45*).

By 24 hours, TTX and BIC produced 9 and 11 altered PiSCES, respectively, most of which were consistent with the trend of upscaling-increased and downscaling-decreased. By 48 hours, 9 additional PiSCES were significant for TTX, and 2 additional PiSCES were significant for BIC. NMDA receptor and Fyn kinase interactions dominated these later-significant PiSCES, perhaps suggesting a delayed weakening/strengthening of NMDARs subsequent to -- and opposite in direction of -- the early AMPAR and mGluR5 tuning. These delayed NMDAR_Fyn (and mGluR5_Fyn) interactions could reflect delayed metaplasticity, a form of homeostatic plasticity distinct from synaptic scaling which adjusts the threshold for LTP and in which Fyn plays a prominent role (*46*, *47*).

### Sensory deprivation induces a homeostatic response in the barrel cortex

We next asked how homeostatic plasticity in vivo compares with synaptic scaling in vitro: do the same sets of PiSCES that change during synaptic scaling also change during prolonged activity manipulation in vivo?

Extreme alteration of activity using sensory deprivation and/or stimulation has been shown to induce homeostatic plasticity in vivo (*3*, *10*, *48*–*52*). Sensory deprivation of the visual cortex, for example, reduces cortical inhibition, which increases spontaneous activity and lowers the threshold for LTP (*48*, *53*–*55*). Similarly, whisker trimming, an established model for sensory deprivation of somatosensory (barrel) cortex (*56*–*58*), has been shown to increase the ratio of excitation to inhibition (E/I) (*57*, *59*). Given this evidence that sensory deprivation alters E/I balance, we asked whether whisker trimming induces an upscaling or downscaling-like homeostatic PIN response.

To test whether we could induce a homeostatic response in vivo, we trimmed the whiskers on one side of an adult mouse (age 7-9 weeks) (**fig. 3A**). After 48 hours, we investigated the functional consequences of whisker trimming by measuring miniature excitatory postsynaptic potentials (mEPSCs) from Layer II/III (L2/3) pyramidal neurons of barrel cortex. The hemisphere ipsilateral to the whisker trim (control side) exhibited a mean frequency and amplitude similar to other reports on WT L2/3 pyramidal cells (**fig. 3B-F**; frequency: 1.87 ± 0.42 Hz; Amplitude: −8.22 ± 2.83 pA) (*60*). Neurons from the contralateral hemisphere (trimmed side), however, demonstrated a 2-fold increase in mean frequency compared with the ipsilateral hemisphere (**fig. 3B,F**; 3.85 ± 0.59 Hz; Student’s t-test, *P* = 0.0035) with no change in mean mEPSC amplitude (**fig. 3C,F**; −6.19 ± 1.18 pA). Cumulative analysis of inter-event intervals revealed a global trend toward decreased time between events onto neurons from the contralateral hemisphere compared with those from the ipsilateral hemisphere (**fig. 3D** Kolmogorov-Smirnov, *P*=0.0242), consistent with an increase in mean mEPSC frequency. Moreover, the cumulative probability of mEPSC amplitude was skewed, revealing a trend toward smaller events that was not reflected in the mean amplitude (**fig. 3E vs. 3C**). There was no difference in spine density between ipsilateral and contralateral hemispheres (**fig. 3G;** 17.25 +/−2.44 vs. 17.13 +/− 1.86 spines/10μm). Taken together, our data confirm that whisker deprivation induced a homeostatic response in the contralateral hemisphere 48 hours after trimming, as previously reported (*56*, *57*, *59*, *61*).

**Fig. 3.**
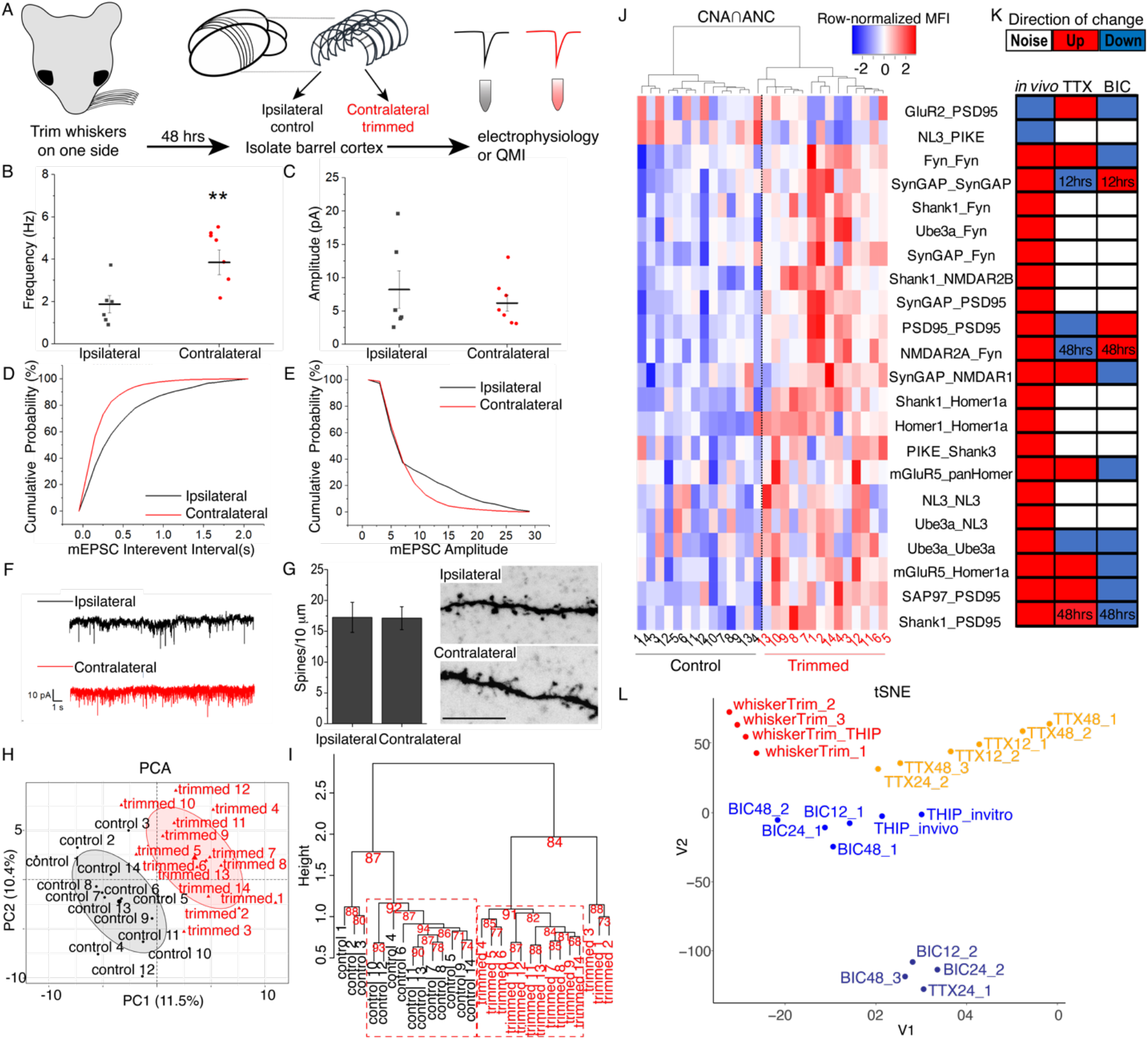
Unilateral whisker trimming causes a homeostatic response in L2/3 pyramidal neurons of the contralateral barrel cortex. **(A)** Workflow of sensory deprivation and analysis of the barrel cortex. **(B,C)** Mean frequency (B) and amplitude (C) of mEPSCs in the ipsilateral (black) and contralateral (red) barrel cortex. **(D)** Cumulative probability of inter-event intervals (0-2s, 0.1s bins). **(E)** Cumulative probability of mEPSC amplitudes (0-30pA, 2pA bins). **(F)** 20s raw traces from ipsilateral (top, black) and contralateral (bottom, red) hemispheres. Scale bar: 10pA, 2s. **(G)** Mean synaptic spine density at 100-150 μm from the soma. Confocal images of neurobiotin-filled L2/3 ipsilateral (top) and contralateral (bottom) dendrites. Ipsilateral: *N* = 9, Contralateral: *N* = 12. Scale bar = 10 μm. **(H)** PCA of all PiSCES (mean MFI > 100) from 14 control (ipsilateral, black) and 14 trimmed (contralateral, red) hemispheres. **(I)** Hierarchical clustering of 14 control and 14 trimmed hemispheres on all PiSCES (mean MFI > 100). AU *P*-values shown in red. **(J)** Heatmap of row-normalized MFI of all high-confidence in vivo PiSCES in control and trimmed samples (N=14 per condition). Samples on the x-axis are arranged by unsupervised hierarchical clustering. **(K)** Chart comparing the direction of change of high-confidence in vivo PiSCES after trimming with the direction of change (if any) after upscaling and downscaling. Red indicates increased interaction, blue indicates decreased, and white indicates that the PiSCES change was not significant. **(L)** tSNE of in vitro upscaling, in vitro downscaling, in vivo whisker trimming, and THIP treatment using mean log_2_(fold change) of all PiSCES significantly altered after TTX, BIC, THIP, or whisker trimming. Each sample input into tSNE represents a single set (N=4 biological replicates per set).

To identify differences in PiSCES between trimmed and control sides, we micro-dissected the barrel cortex from five coronal slices and separated the hemispheres into contralateral (“trimmed”) and ipsilateral (“control”) sides, homogenized the tissue in lysis buffer, and measured PiSCES using QMI (**fig. 3A**) (N=14). PCA and hierarchical clustering of all interactions (MFI>100) showed separation of control and trimmed sides (**fig. 3H,I**). AU *P*-values identified two significant clusters associated with condition (11 samples in each cluster) (**fig. 3I**).

To identify modules of PiSCES significantly correlated with whisker trimming, we performed CNA of all 28 samples (14 in each condition) and intersected with ANC-significant PiSCES. CNA identified two modules (“turquoise” and “green”) that were significantly correlated with whisker trimming**. (fig. S3A, B)** and contained 14 and 8 ANC-significant PiSCES, respectively, that showed clear segregation of conditions when plotted by heatmap **(fig. S3C)**. Together, these 22 ANC∩CNA PiSCES represented high-confidence interactions that changed in response to whisker trimming (**fig. 3J**). These included AMPAR and mGluR interactions, similar to what we observed during in vitro synaptic scaling, including GluR2_PSD95, which was decreased, and mGluR5_Homer and mGluR5_Homer1a, which were increased. Rearrangements in synaptic scaffold proteins were also apparent with PSD95-, SAP97-, Shank1- and Homer-containing interactions uniformly increasing. SynGAP, Shank, and Fyn increased their association with NMDARs, implying increased NMDAR tone and/or decreased threshold for LTP (*46*). Additionally, increased Ube3a interactions implied active protein turnover at the synapse. Overall, these increased interactions suggest broad consolidation of synaptic proteins in effort to increase synapse strength and/or adjust the threshold for LTP.

We next asked how similar these in vivo changes were with in vitro synaptic scaling. Unsupervised hierarchical clustering on these 22 PiSCES separated control and trimmed samples and revealed that most PiSCES increased after trimming (**fig. 3J**). Eleven (50%) were significant at some point during the in vitro time course described above. The direction of the change in vivo, however, was consistent with TTX treatment 6 out of 11 times and consistent with BIC treatment 4 out of 11 times (**fig. 3K**), suggesting that sensory deprivation induces a homeostatic response distinct from in vitro synaptic scaling.

To further examine the relationship between whisker trimming, upscaling and downscaling, we used t-distributed Stochastic Neighbor Embedding (tSNE) of mean log_2_(fold change) values (*62*) and visualized clustering in 2 dimensions **(fig. 3L)**. We included seven TTX experiments (N=4 biological replicates per set) over three time points, seven BIC experiments over three time points, and three whisker trimming experiments. In addition, in order to control for inherent in vitro versus in vivo differences, we included three more sets of experiments that treated in vitro and in vivo samples with the same GABA(A) receptor agonist, THIP **(fig. S4)**. In vivo, THIP (10mg/kg IP) was compared with PBS alone (vehicle control, IP), PBS+whisker trimming, and THIP+whisker trimming after 48 hours of treatment **(fig. S4A)** (*48*, *63*). In vitro, THIP (20μM) was compared with TTX and BIC after 48 hours of treatment **(fig. S4B)**. THIP treatment had a unique effect compared with whisker trimming in vivo and TTX/BIC in vitro and therefore served as a useful in vitro versus in vivo control.

A total of 20 experiments were used as input into tSNE. tSNE of mean log_2_ fold changes of all high confidence PiSCES across all 20 sets revealed that all TTX samples formed one isolated cluster, with the exception of one 24-hour set. The BIC sets formed two clusters, the larger of which contained both the in vitro and in vivo THIP sets. Critically, all four whisker trimming sets formed one independent cluster **(fig. 3L)**, further suggesting that whisker trimming causes a homeostatic response distinct from synaptic upscaling and downscaling.

To determine which PiSCES most contributed to the differences between whisker trimming, TTX treatment, and BIC treatment, we performed pairwise Wilcoxon rank-sum tests of all PiSCES. PiSCES ranked as having the most significant pairwise differences are shown in **table S1** and illustrated in **fig. 4**. Overall, the most significant differences were between the TTX and BIC groups, highlighting the bidirectional nature of upscaling versus downscaling. In addition, several PiSCES were significantly different between whisker trimming and both in vitro groups, including Homer1_mGluR5 (changed in vitro but not in vivo), mGlur5_Fyn (higher magnitude of change in vivo versus in vitro), and Ube3a_Ube3a (deceased in vitro but increased in vivo) **(fig. 4)**. The groups of PiSCES that changed bidirectionally and were most different between TTX and BIC treatment indicated bidirectional shuttling of glutamate receptors via changes in scaffold protein complexes, particularly changes in SAPAP tethering of the Homer-Shank scaffold, as described above. Adjustments of mGluR5 tone via Homer1 and Homer1a, and PI3K signaling, also appeared to play a prominent role in synaptic scaling in vitro **(fig. 4)**. In vivo, many groups of PiSCES behaved similarly to TTX, with increased mGluR5 tone via Homer, increased NMDAR tone via SynGAP, and increased scaffolding interactions, including SAP97_SAPAP and SAP97_PSD95. While the direction of these PiSCES changes may suggest that sensory deprivation in vivo had a similar effect to homeostatic synaptic upscaling, there were additional PiSCES that mirrored downscaling or only changed with sensory deprivation, including Shank1_Fyn, Ube3a_Fyn, and SynGAP_PSD95, emphasizing a unique role for Shank-Fyn-Ube3a complexes and SynGAP signaling in vivo.

**Fig. 4.**
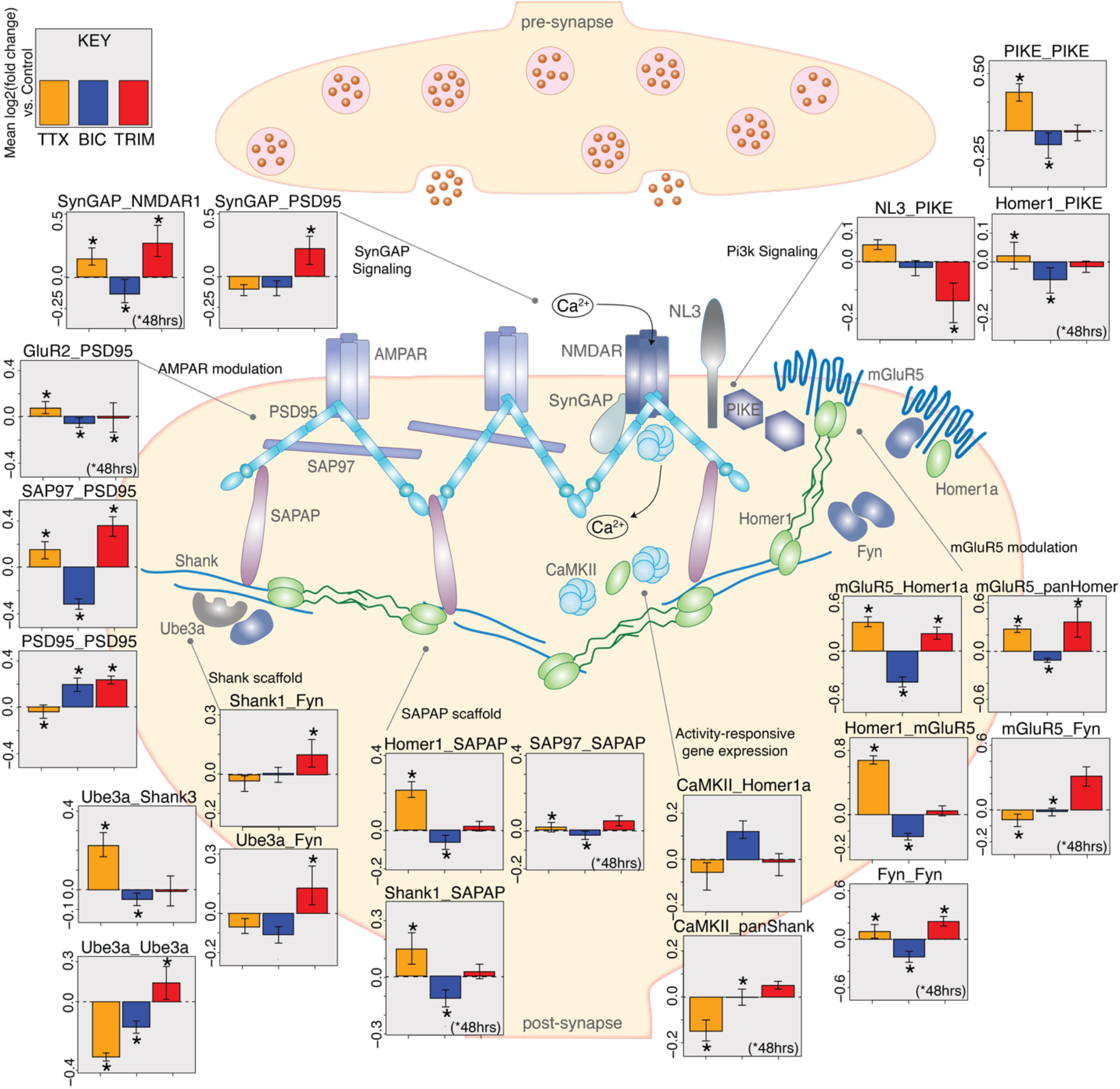
Protein complexes involved in post-synaptic function that change during homeostatic plasticity. Simplified illustration of the excitatory synapse and select functional protein groups evaluated by QMI in the current study. Bar graphs represent the mean log_2_(fold change) over control for the given PiSCES in all sets of experiments using the indicated perturbation. N=6 sets for TTX (outlier in Fig. 3L removed), 7 sets for BIC, and 3 sets for whisker trimming, as shown in Fig. 3L. Error bars represent standard error of the mean. Asterisks indicate that the PiSCES changed significantly in the given condition (ANC∩CNA by 48 hours of treatment or whisker deprivation). All time points are represented, but PiSCES that were not significant until 48 hours of treatment in vitro are indicated (*48hrs).

### Homer1 and Shank3 are synaptic hubs for homeostatic plasticity

We have demonstrated that many interactions involving the synaptic scaffolding protein Homer1 and its short dominant-negative isoform Homer1a are opposite for upscaling and downscaling. In fact, Homer1a is required for downscaling during sleep (*29*), and four PiSCES involving Homer1 and/or Homer1a increased significantly after whisker trimming. We therefore asked whether Homer1 is required for synaptic PIN rearrangements after whisker trimming. We trimmed the whiskers on one side of mice lacking exon 2 of *Homer1*, which abolishes expression of both short and long isoforms of *Homer1* (*64*). We then measured PiSCES by QMI. PCA of all PiSCES (MFI>100) of all four conditions (WT trimmed, WT control, KO trimmed, KO control) revealed that WT and KO samples, regardless of trimming, separated along PC1, while WT trimmed and control samples separated along PC3 (**fig. 5A**). KO trimmed and control samples, however, showed no clear separation.

**Fig. 5.**
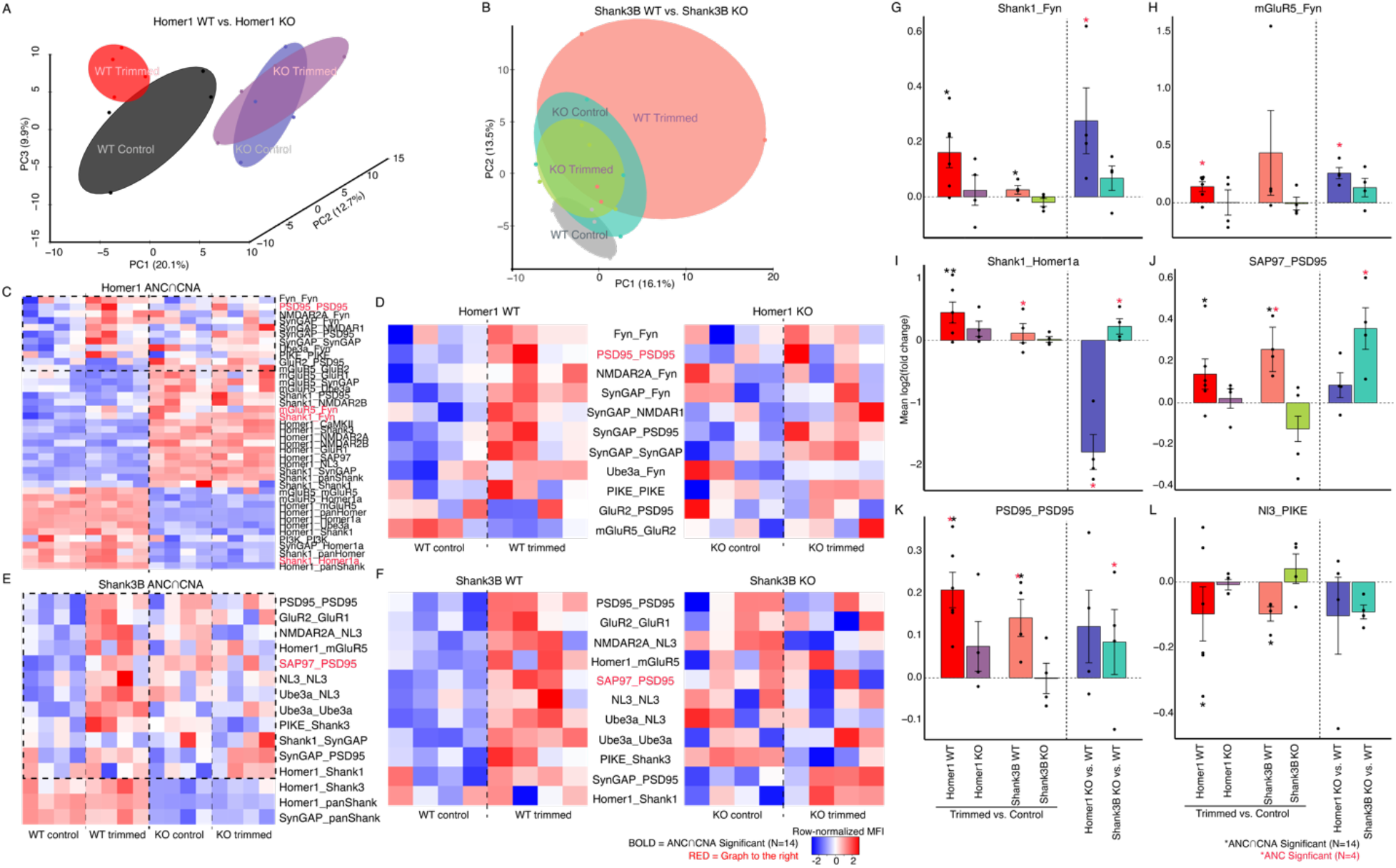
Homeostatic plasticity is disrupted in the barrel cortex of mice mutant for *Homer1* or *Shank3*. **(A, B)** PCA of all PiSCES (mean MFI>100) from 4 biological replicates from sensory-deprived and control hemispheres of *Homer1* WT (red/trimmed vs. black/not trimmed) and *Homer1* KO (purple/trimmed vs. blue/not trimmed) (A) or *Shank3B* WT (grey/trimmed vs. salmon/not trimmed) and *Shank3B* KO (green/trimmed vs. turquoise/not trimmed) (B) adult mice. **(C, E)** Heatmap of row-normalized MFI of all ANC-significant PiSCES after whisker trimming or between WT and KO controls for the *Homer1* set (C) and the *Shank3B* set (E). **(D, F)** Heatmaps of row-normalized MFI of all ANC-significant PiSCES altered only after whisker trimming in WT or KO mice separated into WT and KO specific heatmaps for the *Homer1* set (D) and the *Shank3B* set (F). **(G-L)** Bar graphs of mean log_2_(fold changes) of select PiSCES after whisker trimming in control and KO barrel cortex (red/*Homer1* WT vs. purple/*Homer1* KO; salmon/*Shank3B* WT vs. green/*Shank3B* KO, N=6,4,4,4, respectively) along with baseline differences between WT and KO controls (blue/*Homer1* KO vs. WT and turquoise/*Shank3B* KO vs. WT, N=4,4). The *Homer1* set is on the left, the *Shank3B* set is in the middle, and baseline genotypic differences are on the right. A red asterisk indicates that the PiSCES was ANC-significant for that condition. A black asterisk indicates that the PiSCES was ANC∩CAN-significant for the full WT set (N=14). For C-F, PiSCES that are ANC∩CNA-significant (N=14) are in bold. PiSCES in red are graphed in G-L.

To visualize interactions that were significantly different between trimmed and control sides in both WT and KO barrel cortex, we plotted by heatmap all PiSCES that were significantly altered according to one or more of three pairwise comparisons: 1) WT control vs. WT trimmed, 2) KO control vs. KO trimmed, and 3) WT control vs. KO control (**fig. 5C,D**). Three PiSCES that were elevated after WT whisker trimming (Shank1_NMDAR2B, Shank1_Fyn (**fig. 5G**), and mGluR5_Fyn (**fig. 5H**)) were already elevated in the *Homer1* KO barrel cortex without whisker trimming. Conversely, Shank1_Homer1a (**fig. 5I**), which increased after whisker trimming in WT mice, was absent from the *Homer1* KO barrel cortex. When the MFIs of PiSCES that changed significantly after whisker trimming (**dashed box in fig. 5C**) were plotted in separate heatmaps for WT and KO samples, a clear trend was observed: WT trimmed and control hemispheres, but not KO trimmed and control hemispheres, could be distinguished (**fig. 5D**). Two PiSCES involving SynGAP (SynGAP_SynGAP and SynGAP_PSD95), however, increased after whisker trimming in both WT and KO barrel cortex, indicating that the system did register a change in sensory input but was unable to respond. Together, these findings suggest that whisker trimming had a minimal effect on PIN rearrangements in the sensory-deprived barrel cortex of mice lacking *Homer1*.

Homer1 may drive homeostasis through its interactions with other synaptic scaffolding proteins, such as the Shank family of proteins. Human *SHANK3* is disrupted in an ASD-associated syndrome called Phelan-McDermid Syndrome (*65*, *66*) and has been suggested to play a role in homeostatic plasticity in vitro and firing rate recovery in the mouse visual cortex after visual deprivation (*18*). To determine if *Shank3*, like *Homer1*, is required for PIN rearrangements observed after whisker trimming, we trimmed the whiskers on one side of *Shank3B* KO mice (*67*) and measured PiSCES by QMI. PCA of all PiSCES (MFI>100) of all four conditions revealed that WT trimmed and control samples separated along PC1 and PC2, while KO trimmed and control samples showed no clear separation and overlapped substantially with the WT trimmed group (**fig. 5B**). We plotted all significantly altered PiSCES from the three pairwise comparisons described above by heatmap (**fig. 5E,F**). Like PiSCES in the *Homer1* KO barrel cortex, many of the PiSCES that were elevated after whisker trimming in WT controls were already elevated in the *Shank3B* KO barrel cortex without whisker trimming, including SAP97_PSD95 (**fig. 5J**), PSD95_PSD95 (**fig. 5K**), and NMDAR2A_NL3. Conversely, NL3_PIKE, the only PiSCES that consistently decreased after whisker trimming in the WT barrel cortex, was reduced in the untrimmed KO cortex and did not decrease further after trimming (**fig. 5L**). Homer1_Shank3 increased after whisker trimming in WT mice but was absent in the *Shank3* KO barrel cortex.

When the MFIs of PiSCES that changed significantly after whisker trimming (**dashed box in fig. 5E**) were plotted in separate heatmaps for WT and KO samples, WT trimmed and control sides but not KO trimmed and control sides could be distinguished, consistent with what was observed in *Homer1* KOs (**fig. 5F**). Moreover, SynGAP_PSD95 increased after trimming regardless of genotype, also similar to *Homer1* KOs. Collectively, these results suggest that synaptic scaffold proteins are integral to the ability of the barrel cortex to maintain homeostasis in response to prolonged sensory deprivation, and mice lacking the scaffold proteins Homer1 and Shank3 do not undergo the same PIN rearrangements as WT controls.

## Discussion

Here we have established a “ground truth” for synaptic protein interactions that transduce prolonged activity perturbations to a homeostatic response. We have discovered that prolonged increased or decreased activity of cultured cortical neurons causes widespread bidirectional changes in synaptic PINs that reflect tuning of synaptic strength via insertion/removal of AMPA receptors, remodeling of the Homer-Shank scaffold, and adjustments to mGluR5 tone. Many of these changes appeared stable within as little as 12 hours and persisted throughout the 48-hour time course. Late changes involved Fyn associations with glutamate receptors, which suggested delayed onset of metaplasticity, or delayed adjustments to the threshold for LTP. We also discovered that sensory deprivation induced broad recruitment and consolidation of synaptic proteins that increase mGluR5 tone and increase synaptic transmission and/or lower the threshold for LTP, suggesting a prominent role for metaplasticity in vivo. Finally, we found that this adjustment of synaptic strength upon sensory deprivation via protein rearrangements was abolished in the absence of the scaffold proteins Homer1 or Shank3, despite recruitment of SynGAP signaling, illustrating a prominent role for scaffold proteins in homeostatic plasticity.

### Signal versus noise in signal transduction experiments

QMI attempts to measure unstable protein-protein interactions within noisy and stochastic intracellular environments and thus faces reproducibility challenges similar to mass spectrometry (*68*, *69*). While noise is inherent to all signal transduction experiments, we were able to differentiate signal from noise using CNA modules, which identify PiSCES that change together but may or may not be true positives. For example, while the “turquoise” module shown in figure 1I reveals many true positives, the “brown” module, as shown in figure S2, identifies other PiSCES that change together as a unit but do not correlate strongly with experimental manipulation, implying an underlying biological origin, such as developmental or circadian processes. By using two statistical tests with independent assumptions (ANC and CNA) and carefully matching conditions in each experiment, we can reveal which ANC hits occur together in a module associated with treatment and which occur together in a module associated with noise.

### Bidirectional dynamics of upscaling versus downscaling over time

Homeostatic scaling operates bidirectionally to stabilize cell-wide synaptic strength in response to changes in spiking activity (reviewed in (*7*)) and/or glutamatergic transmission (*2*), reflected, in part, in bidirectional changes in the protein content of the postsynaptic density (*23*). Indeed, it has been established that both upscaling and downscaling employ rapid and delayed protein synthesis (*27*, *28*), and 4 hours of activity blockade was sufficient to employ transcription-dependent processes during upscaling (*20*). In fact, blocking transcription by reducing intracellular calcium with NiCl_2_ or nifedipine was sufficient to produce an upscaling-like response within 24 hours independent of the neuron’s ability to fire action potentials, but the effect was only partial (*20*). Here, we have established that whatever changes in synaptic composition occur during homeostatic scaling, it is how synaptic proteins interact over 48 hours that tunes synaptic strength up or down. We report that, similar to protein turnover, synaptic PINs rearrange bidirectionally with upscaling and downscaling within as little as 12 hours of activity perturbation; however, while most of these early changes were stable after 12 hours, almost half of the significant changes for upscaling were not observed until 48 hours. Collectively, PCA and CNA analyses of the time course data did not indicate a biphasic or multi-step process over time, but rather implied a gradual increase in the intensity of changes over time. The individual PiSCES data, however, told a more sequential story: Changes in AMPAR and mGluR interactions with scaffold proteins occurred early and persisted. Changes in the interaction between the E3 ubiquitin ligase Ube3a with Shanks occurred at intermediate timepoints. By 48 hours, the underlying Shank-PSD cytoskeleton had re-arranged along with changes in NMDAR interactions. Specifically, Fyn interactions with the NMDA receptors 2A and 2B decreased during upscaling and increased during downscaling only at 48 hours, reflecting Fyn adjustment of the threshold for NMDAR-mediated LTP in response to prolonged activity perturbation (*47*). Collectively, these data illuminate a process of rapid AMPAR and mGluR shuttling, driven by SAPAP and Homer1 alterations, followed by a period of ubiquitination of Shank scaffolds, and finally consolidation of PSD95-Shank and NMDAR-Fyn interactions into a new ground state.

### Homeostatic plasticity in vitro versus in vivo

Homeostatic synaptic scaling was first demonstrated in neurons cultured from rat visual cortex (*1*). It has since been demonstrated in vivo in the visual cortex (*10*, *49*, *70*–*72*) and during sleep (*29*). A recent study demonstrated that visual deprivation by eyelid suture elevates spontaneous activity and lowers the threshold for LTP through metaplasticity, or the sliding threshold model (*48*). Similarly, homeostatic plasticity in the barrel cortex after sensory deprivation or GABA agonism is thought to occur not through homeostatic synaptic scaling but through unequal changes in network excitation and inhibition leading to net increased excitation (*59*, *73*). Silencing input to the barrel cortex, therefore, could theoretically result in a counterintuitive elevation in activity and downscaling-like response opposite the response to activity blockade in vitro. While excitatory blockade in vitro was reported to have no effect on excitatory drive onto inhibitory neurons (*1*), inhibitory neurons nonetheless exhibited increased intrinsic excitability after activity blockade in vivo (*74*). Moreover, homeostatic plasticity in inhibitory circuits in vivo appeared to be mediated by specific inhibitory cell types, adding to the vast complexity of in vivo mechanisms that maintain firing rates compared with in vitro (*75*).

Building on the above studies, we observed that in vivo changes to synaptic PINs in response to sensory deprivation exhibit features of both upscaling and downscaling, but do not exactly mimic synaptic scaling. This is consistent with the model that homeostatic mechanisms in vivo involve both excitatory and inhibitory circuits. Like activity blockade in vitro, whisker deprivation resulted in increased mGluR5-Homer interactions, indicating strengthening of mGluR5 tone. Conversely, the prominence of Shank, Fyn, and NMDA receptors in the groups of PiSCES that changed after whisker trimming suggests that metaplasticity, specifically lowered LTP threshold, may be critical in vivo. Nevertheless, we have shown that in all three forms of homeostatic plasticity, the Homer-Shank scaffold plays a central role as a hub for adjusting glutamate receptor strength. Our findings thus far lead to an intriguing hypothesis that different forms of homeostatic plasticity employ different PINs but consistently employ scaffolding protein complexes involving Homers and Shanks.

### Homeostatic plasticity and Autism Spectrum Disorders

Altered E/I has long been hypothesized to be a unifying mechanism for ASD pathology, and several recent reports have suggested that homeostatic plasticity is altered in some ASD syndromes (*16*, *18*). The barrel cortex, which receives tactile input from the whiskers, is a particularly appropriate region for modeling the role of E/I and homeostatic plasticity in ASD, which is often characterized by atypical sensory processing and tactile sensitivity (*76*). One might expect that mouse models of ASD will therefore exhibit altered homeostasis in such a primary sensory area as the barrel cortex (*15*).

Our finding that mice mutant for the ASD genes *Homer1* or *Shank3* fail to exhibit PIN alterations consistent with a homeostatic response to sensory deprivation supports the hypothesis that disrupted homeostatic plasticity, perhaps a result of hyperactive sensory inputs, underlies multiple ASD syndromes. Crucially, our observation that both *Homer1* and *Shank3* KO mice appear to be molecularly “pre-scaled,” or have baseline altered synaptic PINs consistent with sensory deprivation, provides evidence for a “ceiling effect” preventing a homeostatic response to changes in network activity. It is unclear, however, if such a ceiling effect is the result of altered synaptic PINs in excitatory neurons or the result of abnormal circuit development. In other words, *Homer1* and/or *Shank3* KO mice may already compensate for altered E/I balance without sensory perturbation. Indeed, a recent report linked increased firing of excitatory neurons in the vibrissa somatosensory cortex (vS1) of *Shank3B* KO mice with the loss of Shank3 function in *inhibitory* neurons; therefore, inhibitory neurons in *Shank3B* KOs are likely hypoactive, disrupting E/I balance prior to whisker manipulation (*77*). It is also interesting to note that interactions between SynGAP and PSD95 increased after trimming regardless of genotype, suggesting that the established role of SynGAP in gating NMDAR-dependent AMPAR trafficking and synaptic strength via its interactions with PSD95 (*78*, *79*) may operate upstream of scaffolding changes.

Collectively, these results support a model whereby an inability to restore firing rates to a set-point during prolonged activity perturbation is an underlying feature of ASD. Genetic variants that disrupt of any one of these synaptic proteins could alter the neuron’s ability to restore homeostasis during periods of prolonged activity perturbation.

## Materials and Methods

### Animals

All animal work was carried out in compliance with The Seattle Children’s Research Institute IACUC under approved protocol #00072 and institutional and federal guidelines. CD-1 and *Homer1^tm1Mhd^* (stock 023312), and *Shank3^tm2Gfng^* (stock 017688) mice were originally obtained from The Jackson Laboratory (Bar Harbor, ME).

### Genotyping

0.2 μL of crude DNA extract (KAPA Biosystems) from ear punch tissue was used for genotyping the Homer1^tm1Mhd^ allele with the following primers: 5’-CAA TGC ATG CAA TTC CTG AG-3’, 5’-CGA GAA ACT TAC ATA TAT CCG CAA A-3’, and 5’-GAA CTT CGC GCT ATA ACT TCG-3’ (The Jackson Laboratory), or the Shank3B^−^ allele with the following primers: 5’-GAG ACT GAT CAG CGC AGT TG-3’, 5’-TGA CAT AAT CGC TGG CAA AG-3’, 5’-GCT ATA CGA AGT TAT GTC GAC TAG G-3’.

### Cortical Neuron Culture, Drug Treatment, and Surface Labeling

Primary cultures of cortical neurons were prepared as previously described (*37*). Briefly, whole cortex from P0-P1mouse neonates was dissociated using papain (Worthington) and plated at a density of 1.0×10^6^ cells/mL onto 6-well plates treated with poly-D-lysine. Cells were cultured in Neurobasal medium supplemented with 2% B27 and 0.5mM GlutaMax (ThermoFisher) and kept at 37°C, 5% CO_2_ for 18-21 days. After 3-5 DIV, 5-fluoro-2’-deoxyuridine was added to a final concentration of 5 μM to inhibit glial proliferation.

For in vitro homeostatic scaling experiments, TTX (2μM), BIC (40μM), or THIP (20μM) (Tocris) was added directly to the culture medium, and cells were cultured as normal, then lysed after 12, 24, or 48 hours. DMSO (0.4%) served as a vehicle control.

For surface labeling of GluR1, cortical cells were dissociated and cultured as above, with the following exceptions: cells were plated at a density of 1.0×10^5^ cells/mL on glass coverslips treated with poly-D-lysine in 24-well plates. A mouse antibody to the N-terminus of GluR1 conjugated with Alexa Fluor 488 (G-12, Santa Cruz Biotechnology) was added to primary neuron cultures (18-21 DIV) to a final concentration of 4μg/mL (1:50 dilution), and cells were kept at 37°C, 5% CO_2_ for 20 minutes. Culture medium was then removed, and cells were washed once with 1X PBS and fixed with pre-warmed 4% PFA/4% sucrose in 1XPBS at room temperature for 10 minutes shielded from light. After fixation, cells were washed twice with 1X PBS and mounted on glass slides using ProLong Antifade Mountant with Dapi (Invitrogen). Cells were imaged on a Zeiss LSM 710 at 100X. Mean gray values of 8-bit images were calculated in the green channel after thresholding using default settings in ImageJ. Nine cells from at least 3 independent experiments were included in the final analysis. A paired two-tailed Student’s t-Test was used to determine significance.

### Lysate Preparation

Following treatment of cultured cells, cell culture medium was removed and cells were scraped in cold lysis buffer (150mM NaCl, 50mM Tris pH 7.5, 1% NP-40, 10mM Sodium Fluoride, 2mM sodium orthovanadate, protease inhibitor cocktail (Sigma) and phosphatase inhibitor cocktail (Sigma)), transferred to a centrifuge tube, incubated on ice for 15 minutes, and centrifuged at high seed for 15 minutes to remove nuclei and debris. The protein concentration of the supernatant was determined using a Bradford assay (Pierce).

Lysates from cortical slices were prepared as above with the following exceptions: Adult mice were anesthetized with isoflurane and perfused through the heart with ice cold 1X PBS. Brains were immediately removed, placed into cold PBS, and sliced in cold PBS using a VT1200s vibrating microtome (Leica). Five 500μm slices through the somatosensory (barrel) cortex were divided at the midline into ipsilateral and contralateral hemispheres. All slices from one hemisphere were combined into lysis buffer and mechanically dissociated using a PYREX tissue grinder.

### Western Blotting

For western blots, proteins (20 μg per lane) were separated by SDS-PAGE and transferred to a PVDF membrane (Millipore). Membranes were blocked in 4% milk in TBST (0.05M Tris pH7.2, 0.15M NaCl, 0.1% Tween20) for 1 hour at room temperature and incubated with primary antibodies overnight at 4°C or for 1 hour at room temperature. Primary antibodies and dilutions used for western blots were Fyn (clone 59, BioLegend, 1:1000), GluR1 (1504, Millipore, 0.001 mg/ml), Homer1 (AT1F3, LSBio, 1:1000), mGluR5 (5675, Millipore, 1:2000), PSD95 (K28/43, BioLegend, 1:500), beta-Actin (GTX109639, GeneTex, 1:10,000), p-ERK T202/Y204 (4370, Cell Signaling Technologies, 1:1000), and total ERK (9102, Cell Signaling Technologies, 1:1000). Primary antibodies were detected using species-specific HRP-conjugated secondary antibodies. Blots were developed using Femto Maximum Sensitivity Substrate (Pierce) and imaged using a ProteinSimple imaging system (San Jose, CA).

### Quantitative Multiplex Immunoprecipitation

QMI was performed as described previously (*37*, *38*). Briefly, a master mix containing equal numbers of each antibody-coupled Luminex bead was prepared and distributed to lysates containing equal amounts of protein and incubated overnight on a rotator at 4°C. The next day, each sample was washed twice in cold Fly-P buffer (50mM tris pH7.4, 100mM NaCl, 1% bovine serum albumin, and 0.02% sodium azide) and distributed into twice as many wells of a 96-well plate as there were probe antibodies (for technical duplicates). Biotinylated detection (probe) antibodies were added to the appropriate wells and incubated at 4°C with gentle agitation for 1 hour. The resulting bead-probe complexes were washed 3 times with Fly-P buffer, incubated for 30 minutes with streptavidin-PE on ice, washed another 3 times, resuspended in 125μl ice cold Fly-P buffer, and processed for fluorescence using a customized refrigerated Bio-Plex 200 (*38*). Antibody clone names, catalog numbers, and lot numbers are the same as in (*37*).

### Whisker Trimming and THIP administration

Mystacial whiskers on one side of an adult mouse (7-9 weeks) were removed with an electric trimmer while the mouse was under very brief isoflurane anesthesia. For THIP experiments, either THIP hydrochloride (Tocris) or an equal volume of PBS (vehicle control) was injected intraperitoneally at a concentration of 10mg/kg. Eight hours after the initial injection at the time of trimming, a second injection was given, and two additional injections were given the following day, 8 hours apart, for a total of 4 injections over 48 hours. Lysates were prepared as described above 48 hours after trimming.

### Data Analysis and Statistics

#### ANC

High-confidence, statistically significant differences in bead distributions between two conditions for individual PiSCES, after correcting for multiple comparisons, were identified using ANC as described in (*38*). ANC most consistently identifies significant PiSCES changes in N of 4 experiments, becoming progressively more stringent with any N over 4. We therefore divided biological replicates into sets of four or five by date of experiment, so that experiments that were performed consecutively were grouped for ANC analysis. Any PiSCES that was found to be significant by any ANC comparison was considered a “hit”.

#### CNA

Modules of PiSCES that co-varied with experimental conditions were identified using CNA as described in (*37*, *38*). Briefly, bead distributions used in ANC were collapsed into a single MFI value for every PiSCES and averaged across technical replicates for input into the WGCNA package for R (*41*). PiSCES with MFI < 100 were removed, and batch effects were corrected using COMBAT (*80*). Power values giving the approximation of scale-free topology were determined using soft thresholding with a power adjacency function. The minimum module size was always set to between 10 and 12, and modules whose eigenvectors significantly correlated with an experimental trait (p<0.05) were considered “of interest.” PiSCES belonging to a module of interest and whose probability of module membership in that module was < 0.05 were considered significantly correlated with that trait. PiSCES that were significant by both ANC and CNA for a given experimental condition were considered significantly altered in that condition.

#### Hierarchical clustering and PCA

Post-COMBAT, log_2_ transformed MFI values were clustered using the hclust function in R with a correlation distance matrix and average clustering method. Approximately unbiased (AU) values were determined using the pvclust package in R (*40*). PCA was performed using the prcomp function in R.

#### t-SNE

Log_2_(fold change) values for all PiSCES, or for all PiSCES that were significant in any condition where specified, were encoded in a vector for each condition and dimensionally reduced using PCA. The t-SNE algorithm (*62*) with perplexity 2 was then used to reduce the PiSCES vectors to 2 dimensions in R. Hierarchical clustering was used to fit conditions into 4 clusters, which were visualized using ggplot. The most significant PiSCES between treatment groups were determined by Wilcoxon rank-sum pairwise comparisons of sets grouped by treatment, regardless of tSNE cluster, in R. Heatmaps were generated using Heatmap.2 in R.

### Electrophysiology

#### Acute slice preparation

Adult male mice (7-9 weeks old) were anesthetized with isoflurane inhalation anesthetic and perfused through the heart with ice cold standard artificial cerebral spinal fluid (ACSF). Brains were rapidly removed and placed into ice-cold cutting ACSF. 300 μm thalamocortical slices were cut on a VT1000s vibrating microtome (Leica) and were gently warmed to 35⁰C for 15-20 minutes then allowed cooled to room temperature prior to recording.

#### Patch clamp recording

All recordings were performed at 32.5 ± 1⁰C. Barrel cortex LII/III pyramidal neurons were visually identified under DIC optics (Nikon) and confirmed by their electrophysiological profile and *posthoc* morphological reconstruction. Recordings were sampled at 10 KHz with a Multiclamp 700B amplifier Digidata 1400 digitizer (Molecular Devices) and were rejected if I_Holding_ exceeded ± 100 pA from −70 mV in voltage camp or if Vm changed more than 15% in current clamp. Only cells with > 1 GΩ seal and Vm < −55 mV were included in analysis. Junction potential was 13 mV (compensated).

#### Histology

Neurobiotin was allowed to passively diffuse throughout the dendritic tree for the duration of each recording (>10 minutes). The recording pipette was carefully withdrawn and the membrane allowed to re-seal before separation with a tap to the headstage. The slice was immediately transferred to 4% paraformaldehyde for > 1hr and kept up to 2 weeks at 4⁰C before processing with streptavidin. PFA was removed with 4 15-minute washes in 1X PBS and followed with a 2-hr block in 1X PBS + 0.5% Triton X-100 and 10% goat serum. 1% Alexa 488-streptavidin conjugate was added to fresh blocking solution for 1-2 hours then washed 4X for 15 minutes each in 1X PBS. Slices were mounted with Vectashield anti-fade mounting medium with DAPI (Vector Labs) and visualized on a LSM 710 confocal microscope (Zeiss) with 20X, 40X, and 63X objectives. Spines were counted by an observer blind to hemisphere over a 15-50 μm continuous section of dendrites located 100-150 μm from the cell body.

#### Solutions and drugs

Standard ACSF (in mM): NaCl 128, KCl 3, NaH_2_PO_4_ 1.25, NaHCO_3_ 26, Glucose 10, CaCl_2_ 2, MgSO_4_ 2; pH 7.35-7.4 and 305-315 mOsm. Cutting ACSF (in mM): Sucrose 75, NaCl 87, KCl 3, NaH_2_PO_4_ 1.25, NaHCO_3_ 26, Glucose 20, CaCl_2_ 0.5, MgSO_4_ 7; pH 7.35-7.4 and 305-315 mOsm. Internal recording solution (in mM): KmeSO_4_, KCl 7, EGTA 0.1, Na_2_ATP 2, MgATP 2, Na_2_GTP 0.3, phosphocreatine 5, 0.2% neurobiotin. 7.4 pH, 285-295 mOsm. Bath-applied drugs: Tetrodotoxin 1 μM, picrotoxin 25 μM.

#### Analysis and statistics

Raw traces were acquired, offline filtered to 1 KHz, and analyzed with the pClamp software suite (v. 10.7, Molecular Devices). Neuron reconstructions were performed using ShuTu dendrite tracing software (http://personal.psu.edu/dzj2/ShuTu/) and the Sholl Analysis plug-in for FIJI/ImageJ (Ferreira et al. Nat Methods 11, 982-4 (2014)). Statistics and plotting were performed with OriginLab Pro 2017.

## Acknowledgments

The authors would like to thank members of the Smith lab, especially Emily Brown, and members of the Center for Integrative Brain Research for helpful discussions.

## Author contributions

W.E.H. and S.E.P.S designed the study. W.E.H., H.S., J.L., K.I., and E.G. performed the experiments. W.E.H., H.S., J.L., and S.E.P.S. analyzed the data. W.E.H., H.S., and S.E.P.S. wrote the manuscript.

## Supplementary Materials

**Fig. S1.**
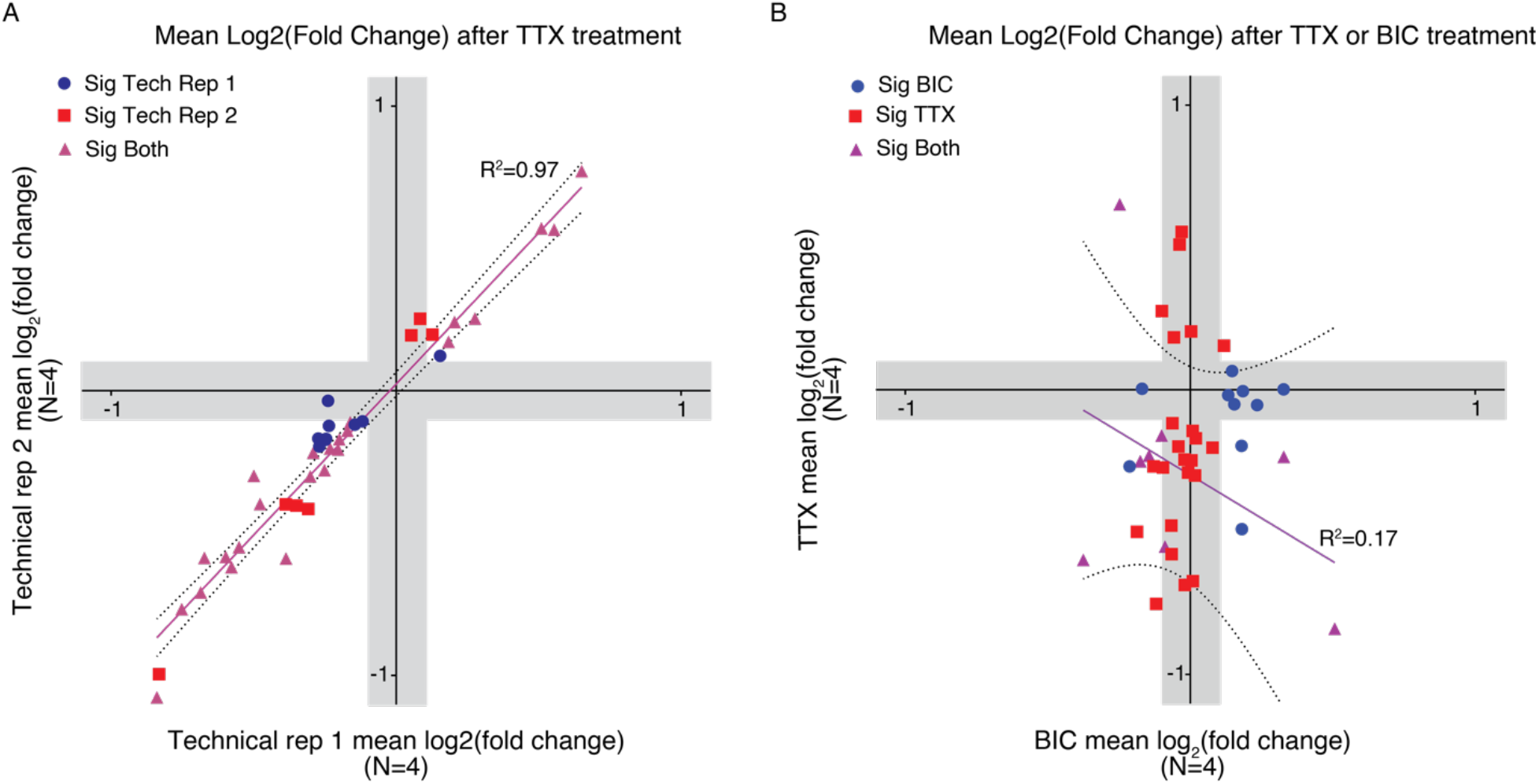
Parallel experiments show high concordance. **(A, B)** Mean log2(fold change) of all PiSCES that changed significantly in at least one of the two sets of experiments being compared. Each set was run in parallel (on the same QMI plate). Sets under a single treatment condition (TTX1 vs. TTX2) showed high concordance (A), while sets run in parallel under different treatment conditions (TTX1 vs. BIC1) showed low concordance (B).

**Fig. S2.**
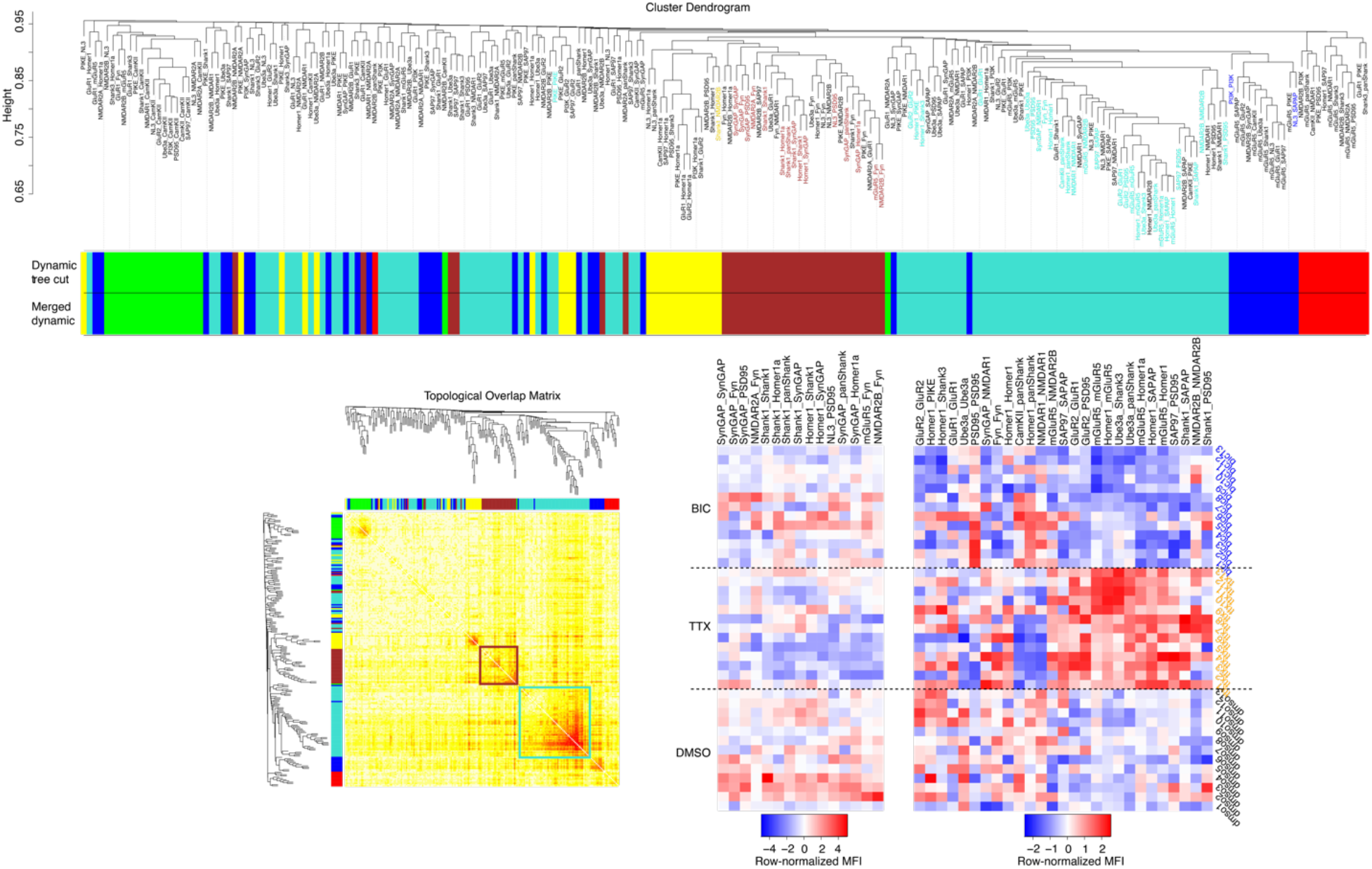
One bidirectional module, distinct from a “noise” module, contains high-confidence PiSCES that change after 48 hours of upscaling or downscaling. From bottom left, clockwise: Topological overlap matrix (TOM) plot (left) and dendrogram (top) of all PiSCES (MFI>100) with the color of the primary module of each indicated below the dendrogram. Heatmaps (bottom right) of row-normalized MFIs of all ANC-significant PiSCES in the brown module (left heatmap) or turquoise module (right heatmap). Samples on the Y-axis are arranged by treatment, and PiSCES on the X-axis are arranged in the order they appear in the dendrogram.

**Fig. S3.**
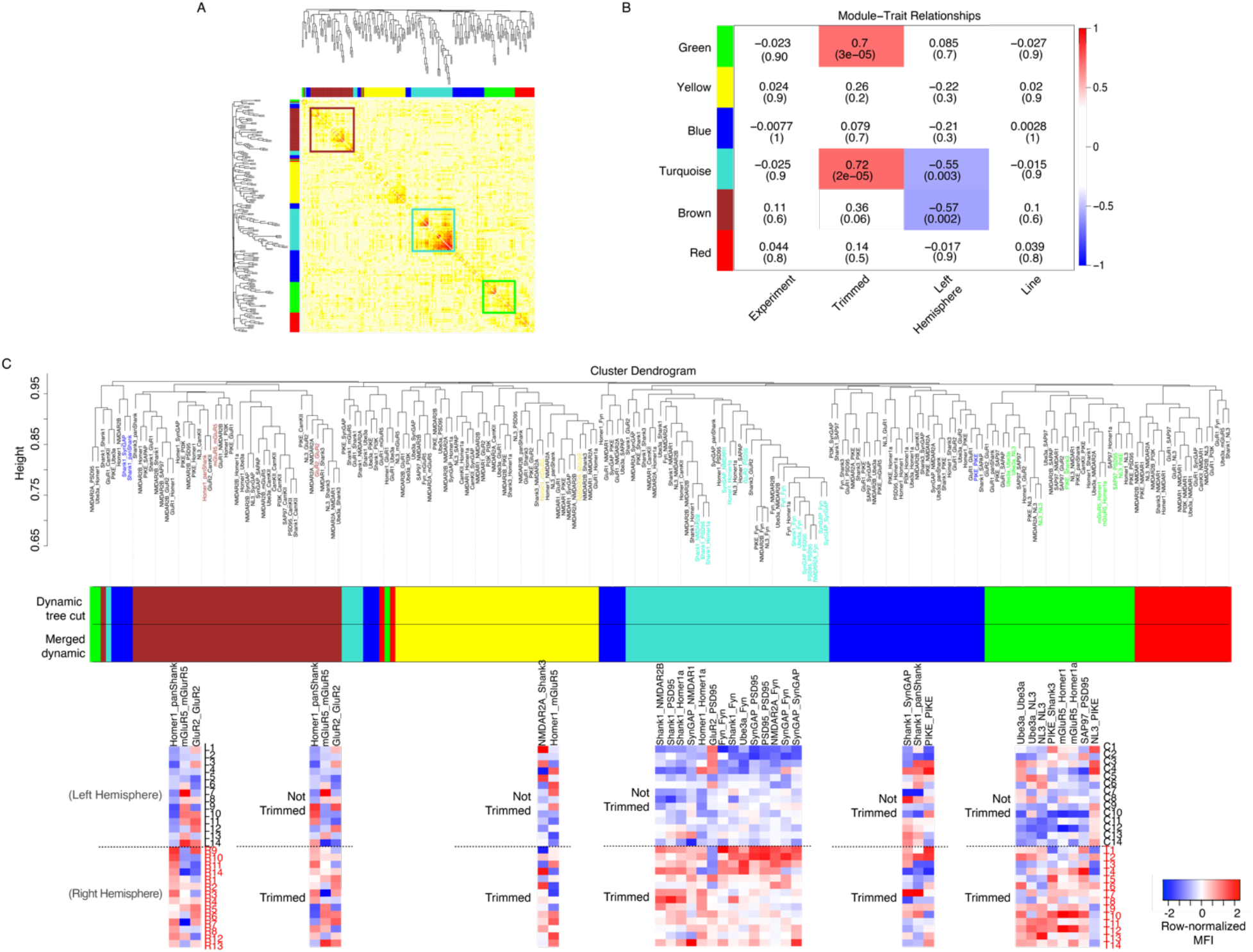
QMI detects a subset of high-confidence PiSCES changes during *in vivo* homeostatic plasticity. **(A)** Topological overlap matrix (TOM) plot of all PiSCES (MFI>100) with and without whisker trimming. **(B)** Module-trait relationship heatmap showing the correlation (top number) and *P*-value for each module-trait pair (N=14 biological replicates). **(C)** TOM-clustering dendrogram of PiSCES arranged above their primary modules, with heatmaps of row-normalized MFIs of all ANC-significant PiSCES in the brown (far left), yellow (left of center), turquoise (center), blue (right of center), or green (far right) module. Samples on the Y-axis are arranged by treatment except for the far left heatmap, in which they are arranged by hemisphere. PiSCES on the X-axis are arranged in the order they appear in the dendrogram. Abbreviations: Control (C), Trimmed (T), Left hemisphere (L), and Right hemisphere (R).

**Fig. S4.**
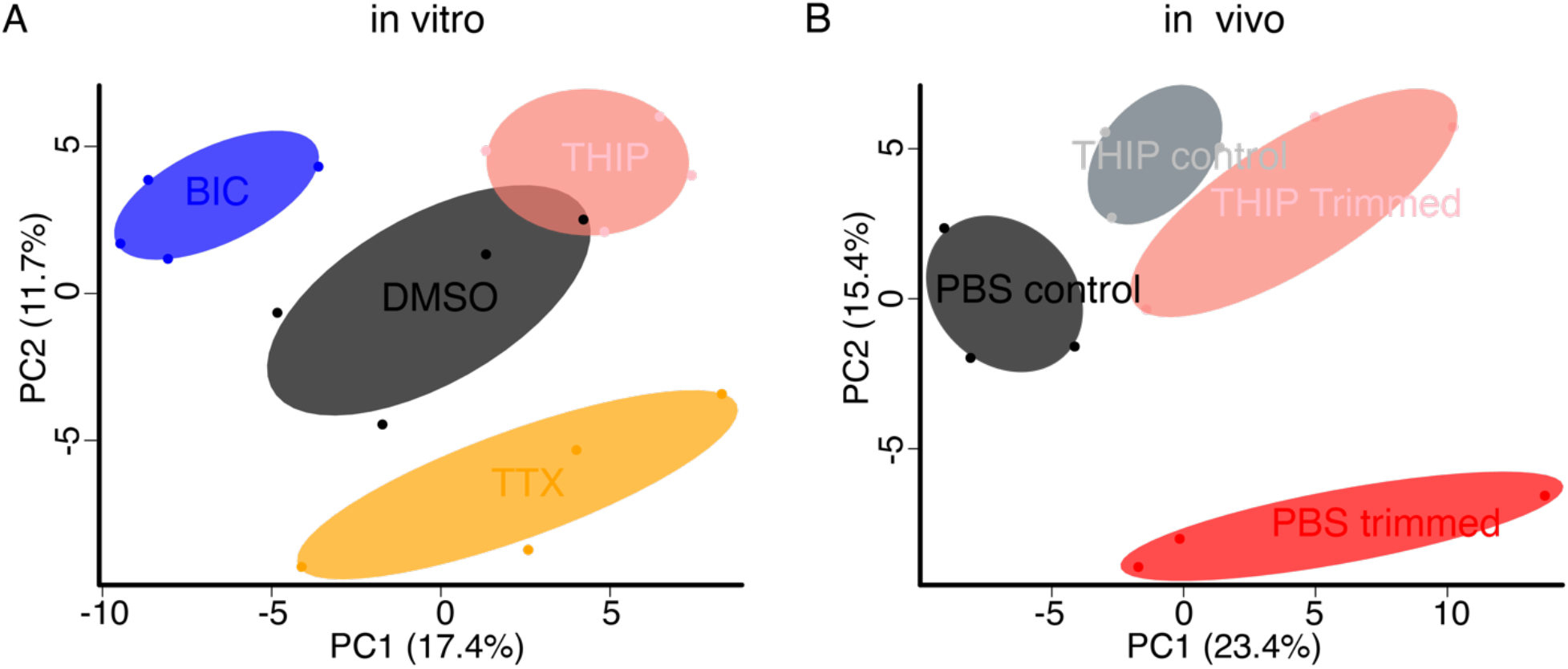
THIP treatment causes synaptic protein rearrangements that are distinct from in vitro homeostatic synaptic scaling and in vivo whisker deprivation. **(A)** PCA of all PiSCES (MFI>100) after 48 hours of DMSO (control, grey), TTX (orange), BIC (blue), or THIP (pink) treatment of cultured cortical neurons. **(B)** PCA of all PiSCES (MFI>100) in the barrel cortex after 48 hours of PBS alone (grey), PBS plus whisker trimming (red), THIP alone (light grey), or THIP plus whisker trimming (pink).

**Table S1.** Top-ranked significantly different PiSCES (by Wilcoxon rank-sum test) between three models of homeostatic plasticity. PiSCES in bold are significantly different between the in vivo model and both in vitro models. N=7 sets (TTX), 7 sets (BIC), and 4 sets (Trim) with 4 biological replicates per set.

| TTX vs. BIC | *P*-value | TTX vs. Trim | *P*-value | BIC vs. Trim | *P*-value |
| --- | --- | --- | --- | --- | --- |
| CaMKII_PIKE | 0.003 | CamKII_panShank | 0.028 | Fyn_Fyn | 0.023 |
| Homer1_mGluR5 | 0.003 | GluR2_GluR1 | 0.028 | **Homer1_mGluR5** | 0.023 |
| mGluR5_panHomer | 0.003 | **Homer1_mGluR5** | 0.028 | mGluR5_Homer1a | 0.023 |
| mGluR5_Homer1a | 0.003 | Homer1_SAPAP | 0.028 | SAP97_PSD95 | 0.023 |
| SAP97_PSD95 | 0.003 | **mGluR5_Fyn** | 0.028 | Ube3a_Fyn | 0.023 |
| Ube3a_panShank | 0.003 | NL3_PIKE | 0.028 | **mGluR5_Fyn** | 0.040 |
| Ube3a_Shank3 | 0.003 | SynGAP_PSD95 | 0.028 | mGluR5_panHomer | 0.040 |
| NMDAR1_NMDAR1 | 0.005 | Ube3a_panShank | 0.028 | PIKE_PSD95 | 0.040 |
| Homer1_SAPAP | 0.008 | Ube3a_Shank3 | 0.028 | SynGAP_NMDAR1 | 0.040 |
| NMDAR2B_SAPAP | 0.008 | **Ube3a_Ube3a** | 0.028 | **Ube3a_Ube3a** | 0.040 |
| Shank1_SAPAP | 0.008 |  |  |  |  |
| Ube3a_Ube3a | 0.008 |  |  |  |  |
| Homer1_NMDAR1 | 0.012 |  |  |  |  |
| mGluR5_mGluR5 | 0.012 |  |  |  |  |
| CaMKII_Homer1a | 0.018 |  |  |  |  |
| Homer1_panHomer | 0.018 |  |  |  |  |
| PSD95_PSD95 | 0.018 |  |  |  |  |
| Fyn_Fyn | 0.027 |  |  |  |  |
| Homer1_Shank1 | 0.027 |  |  |  |  |
| PIKE_PIKE | 0.027 |  |  |  |  |

